# Real-Time Cell Analysis Reveals Distinct Roles of S100A4 in Regulating Proliferation, Migration, and Invasion of JAR Choriocarcinoma Cells

**DOI:** 10.64898/2026.02.26.708094

**Authors:** Bin Yu, Hai-Gang Ding, Feng Zhang, Hong-Mei Lin, Gui-Yu Xia, Ye-Jun Jiang, Jian Zhao, Guo-Ping Li, Jin-Long Ding, Na Ding, Xin-Yue Zhang, Hai-Tao Pan, Ping Ying, Yao He

## Abstract

S100A4, a metastasis promoting calcium binding protein, drives tumor progression through pleiotropic mechanisms, yet its context dependent functions in gestational malignancies remain elusive. To dynamically decode its role in choriocarcinoma pathogenesis, we leveraged label free real time cell analysis (RTCA) to profile malignant phenotypes in JAR cells following siRNA mediated S100A4 silencing, complemented by apoptosis assessment and targeted signaling profiling. Efficient knockdown (verified by qPCR/Western blotting) significantly attenuated cellular proliferation (96 hr cell index slope decreased vs. scramble control; p<0.01) and suppressed migration capacity (p<0.01). Critically, S100A4 depletion did not induce apoptosis (flow cytometry and cleaved caspase 3/9 blotting confirmed no significant change), and invasion through Matrigel coated membranes remained statistically unaltered despite comparable experimental rigor. Mechanistically, S100A4 silencing triggered adaptive signaling rewiring: IRS1 and PI3K expression were elevated, Akt1 was suppressed, while MEK1/2 remained unchanged suggesting compensatory pathway activation.

## 1. Background

Choriocarcinoma represents one of the most clinically aggressive gestational malignancies, distinguished by its propensity for early vascular invasion and distant metastasis. Despite advances in chemotherapy, patients with refractory or recurrent disease face mortality rates exceeding 30%^1^, underscoring an urgent need to dissect the molecular mechanisms governing its disseminative traits. Critical among potential drivers is the metastasis-associated calcium-binding protein S100A4^2^, S100A4 is upregulated in various cancers and significantly promotes cancer cell proliferation, migration, invasion, and EMT through the regulation of multiple signaling pathways and molecular mechanisms^3–5^, but remain underexplored in trophoblastic contexts. Its dysregulation in placental pathologies^6^ and association with poor prognosis in epithelial tumors^7^ suggest a plausible, yet unverified, function in choriocarcinoma progression.

Current understanding of S100A4’s oncogenicity, however, relies heavily on endpoint assays that capture static snapshots of cell behavior—such as fixed-timepoint Transwell invasion or colony formation^8^—while overlooking dynamic phenotypic adaptations. These methods cannot resolve kinetic nuances in cellular responses, potentially masking context-dependent functional hierarchies. Moreover, the predominance of correlative evidence from tissue studies fails to establish causal links^9^ between S100A4 expression and specific malignant behaviors. This knowledge gap is particularly consequential for choriocarcinoma, where the unique biology of trophoblastic cells may engender distinct regulatory dependencies compared to carcinomas of epithelial origin^10^.

To address this limitation, we leveraged Real-Time Cell Analysis (RTCA)^11^, an impedance-based platform enabling continuous, label-free monitoring of cellular functions. Unlike conventional approaches, RTCA quantifies phenotypic dynamics—from proliferation kinetics to migration velocity and barrier penetration—at minute-scale resolution under physiologically relevant conditions^12^. While S100A4’s role in proliferation and migration is established, its impact on apoptosis and downstream signaling in trophoblastic malignancies remains ambiguous. To comprehensively dissect S100A4’s oncogenic mechanisms, we integrated apoptosis assessment (via flow cytometry and caspase-3/9 blotting) and targeted signaling profiling (IRS1/PI3K/Akt/MEK axes) alongside RTCA phenotyping. This multi-layered approach addresses whether S100A4 silencing triggers compensatory pathway activation to sustain invasion despite proliferative/migratory suppression. By integrating siRNA-mediated S100A4 silencing with multi-parametric RTCA profiling in human choriocarcinoma JAR cells, this study aims to: (1) delineate the real-time contribution of S100A4 to proliferation, migration, and invasion; (2) test the hypothesis that S100A4 exhibits phenotype-selective regulation^13^ rather than uniform control across malignant hallmarks. Our approach not only circumvents the artifacts of endpoint fixation but also provides unprecedented temporal resolution for functional stratification in trophoblastic malignancies.

## 2. Materials & Methods

### 2.1. Cells and cell culture

JAR cells were maintained in Roswell Park Memorial Institute (RPMI)-1640 medium (Gibco, Cat. no. 21875-042) supplemented with 10% fetal bovine serum (Gibco, Cat. no. 10099141), under humidified conditions at 37°C with 5% CO_2_. When the cells reached 80%–90% confluence, they were digested with 0.25% trypsin-EDTA (Gibco, Cat. no. 25200056) and passaged at a split ratio of 1:2 to maintain them in the logarithmic growth phase for subsequent experiments.

### 2.2. Small interference RNA (siRNA) treatment

Cells were seeded into six-well plates in complete medium for 24 h prior to transfection. Transfection was carried out with Lipofectamine 6000 reagent (Beyotime, Cat. no. C0526) following the manufacturer’s protocol. Cells were transfected with 20 μM control siRNA or target siRNA oligos (Ribo, Cat. no. SIGS0007750-1). Transfected cells were cultured in complete medium at 37°C for 48 h.

### 2.3. Real-time cell analysis (RTCA)

RTCA is a novel label-free analytical technique capable of non-invasively monitoring cellular behaviors, including proliferation, migration, invasion, etc. In this study, we used the xCELLigence DP RTCA instrument from Agilent Technologies to perform relevant experiment.

#### 2.3.1 Proliferation assay

Cells were digested with trypsin for 2 min after transfection, resuspended in complete medium, counted, and adjusted to a concentration of 6×10^4^ cells/mL. 50 μL medium was added to each well of the E-16 culture plate (Agilent, Cat. no.300601010) to acquire background signals. After 30 minutes of incubation in a cell culture incubator, baseline calibration of the plate was performed using the RTCA software. Subsequently, 100 μL of the cell suspension was added to each well to seed the cells, and cell proliferation was monitored until the experiment concluded.

#### 2.3.2 Migration assay

Cell migration was evaluated by RTCA method on a CIM plate (Agilent, Cat. no.05665817001). Briefly, 165 μL of complete medium was added to the lower chamber. After assembling the upper and lower chambers, 30 μL of serum-free medium was added to the upper chamber. The plate was equilibrated at 37°C with 5% CO_2_ for 1 hour, followed by baseline measurement. Then, 100 μL of serum-free cell suspension containing 40,000 cells was added to the upper chamber per well. After allowing the plate to settle at room temperature for 30 minutes, real-time impedance measurements were performed.

#### 2.3.3 Invasion assay

To ensure uniform coating of Matrigel (Corning, Cat no. 354234) in CIM plates, centrifuge tubes and the upper chamber of the CIM-Plate were pre-cooled at 4 °C overnight. Matrigel was thawed at 4 ° C overnight. Matrigel was diluted with serum-free medium at a ratio of 1:60 on ice. To each well of the upper chamber, 50 μL of the diluted Matrigel was added, followed by removal of 30 μL. The plate was then incubated at 37 °C for 4 hours until Matrigel solidified. The remaining procedures were identical to those of the cell migration assay.

### 2.4. Western blot analysis

According to the kit instructions, total protein was extracted from cells with RIPA buffer (Beyotime, cat no. P0013C), and its concentration was determined by the BCA assay (Beyotime, cat no. P0012S). Equal amounts of protein were separated by 10% SDS-PAGE (Sagon, cat. no. C651101), transferred onto a PVDF membrane (Beyotime, cat. no. FFP24), blocked with TBST containing 5% skim milk at room temperature for 1 hour (Beyotime, cat. no. P0233), and subsequently incubated overnight at 4°C with primary antibodies against Caspase-3 (1:1000; Beyotime, cat no. AF1213), Caspase-9 (1:1000; Beyotime, cat no. AF1264), Tublin (1:1000; Beyotime, cat no. AT819), IRS1 (1:500; Beyotime, cat no. AF7299), PI3-K (1:1000; Beyotime, cat no. AF1966), Akt1 (1:1000; Beyotime, cat no. AF0045), MEK1/2 (1:1000; Beyotime, cat no. AF1057), GAPDH (1:1000; Beyotime, cat no. AF5009), S100A4(1:1000; Beyotime, cat no. AF5291), and β-Actin (1:1000; Beyotime, cat no. AF2811). After three washes with TBST, the membrane was incubated with horseradish peroxidase (HRP)-conjugated secondary antibody at room temperature for 1 hour: HRP-labeled Goat Anti-Mouse IgG (1:1000; Beyotime, cat no. A0216) and HRP-labeled Goat Anti-Rabbit IgG (1:1000; Beyotime, cat no. A0208). Protein bands were visualized using an ECL reagent (Beyotime, cat. no. P0018AS), and band intensities were normalized to an internal reference protein for standardization.

### 2.5. RT-qPCR analysis

Total RNA was isolated with TRIzol (Tiangen, cat. no. DP419), then reverse-transcribed into cDNA employing the PrimeScript RT Kit (Tiangen, cat. no. KR116-01) and miRNA-specific stem-loop primers targeting specific miRNAs to enhance specificity. RT-qPCR assays were conducted using SYBR Premix Ex Taq (Tiangen, cat. no. RK145) and the corresponding primers, with normalization to U6 small nuclear RNA. Relative gene expression levels were determined by the 2−ΔΔ Ct method.

### 2.6. Flow Cytometry Analysis

JAR cells were measured for apoptosis levels using the Annexin V-FITC/PI Apoptosis Detection Kit (Yeasen, cat. no. 40302). Two days after transfection of HTR8 cells, the cells were collected by treatment with EDTA-free trypsin(Sagon Biotech, cat. no. E607003), washed with Pre-cooled PBS and treated according to the kit’s instructions. Briefly, 1-5×10^5^ cells were suspended in 100 μL of binding buffer and put in a centrifuge tube, 5 μL Annexin V-FITC and 10 μL PI Staining Solution was added, and then incubated in the dark for 10 minutes. Subsequently, 400 μL of binding buffer was added to the tubes and assayed by flow cytometry. Data were processed using FlowJo v10.6.2 software (Treestar, Ashland, OR, USA).

## 3. Result

First, we aimed to downregulate the mRNA expression of S100A4 in JAR cells using siRNA technology. To this end, JAR cells were transfected with S100A4-specific siRNA sequences and indicated as siS100A4/JAR cells, whereas JAR cells transfected with a scrambled siRNA sequence (serving as a negative control) were appointed as Ctrl/JAR cells. RT-qPCR analysis confirmed a significant reduction in S100A4 mRNA expression in the transfected JAR cells. Western blotting was subsequently performed to evaluate the protein expression of S100A4 in these cells (Fig. 1a). Following siRNA-mediated silencing, the expression of S100A4 protein was markedly decreased (Fig. 1b).

**Fig. 1.**
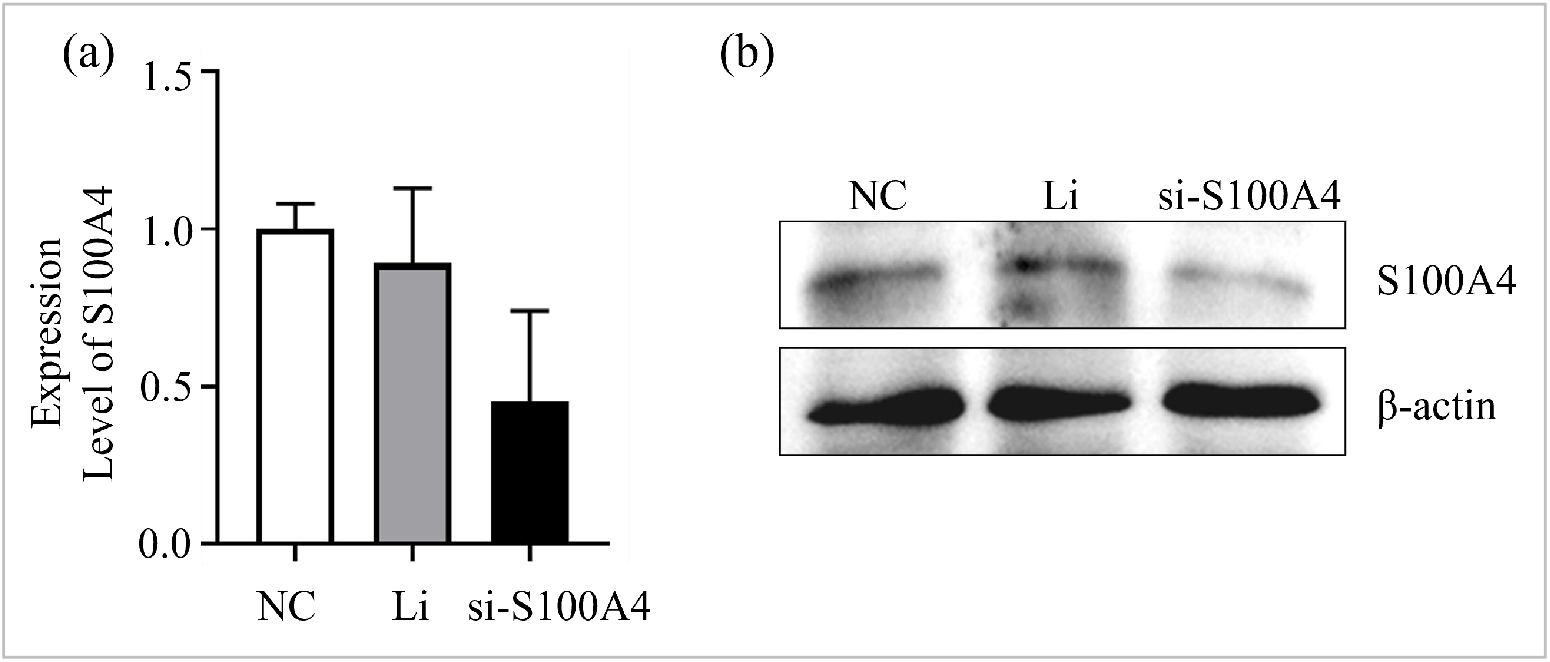
S100A4 expression is decreased in siS100A4 -transfected JAR cells. (a) RT-qPCR analysis of S100A4 expression in siCtrl and siS100A4/ JAR cells. β-actin was used for normalization (b) Western blot analysis of S100A4 expression in siCtrl and siS100A4/ JAR cells. β-actin is used as a loading control.

Real-time cell analysis (RTCA) is a novel technique that employs real-time cellular monitoring to detect the proliferation, migration, and invasion of cells during cell culture. It enables uninterrupted, label-free, and real-time analysis of cells throughout the experimental process. Currently, RTCA is widely applied by researchers worldwide across many diverse research fields. We evaluated the proliferation dynamics of S100A4-knockdown JAR cells using xCELLigence RTCA. Cell index profiles showed that S100A4-siRNA-transfected JAR cells exhibited decreased proliferation capacities (Fig. 2).

**Fig. 2.**
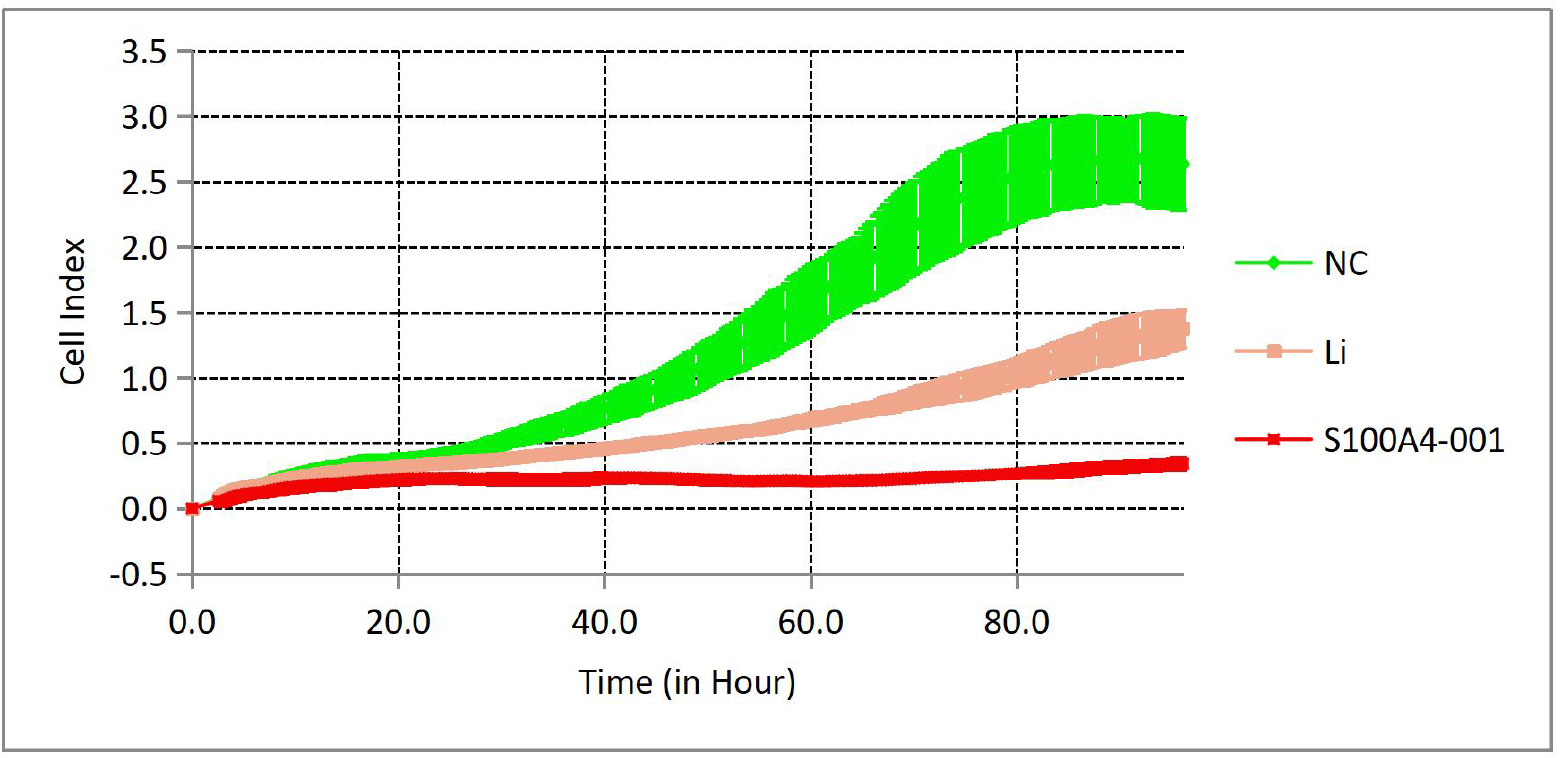
Knockdown of S100A4 inhibited cell proliferation in JAR cell line. Real time cell analysis showing that knockdown of S100A4 inhibited cell proliferation of JAR cells.

Cell migration, a fundamental property of cells, occurs in numerous physiological and pathological processes, wherein migrating cells play a critical role in tissue morphogenesis during development, post-developmental tissue repair, and the support of tumor invasion and metastasis^14^. Similarly, to evaluate the effect of S100A4 knockdown on the migration and invasion capacities of JAR cells, we transfected JAR cells with siRNA-S100A4. Following transfection, both siS100A4-transfected JAR cells and control cells (scrambled siRNA-transfected) were seeded into CIM-Plates. The migration and invasion dynamics of these cells were then monitored in real-time using the RTCA system. The results showed that compared with the control group, knockdown of S100A4 reduced the migration ability of JAR cells (Fig.3), but did not reduce their invasion ability (Experimental results of the liposome reagent control group and the knockdown of S100A4 experimental group were similar, Fig.4).

**Fig. 3.**
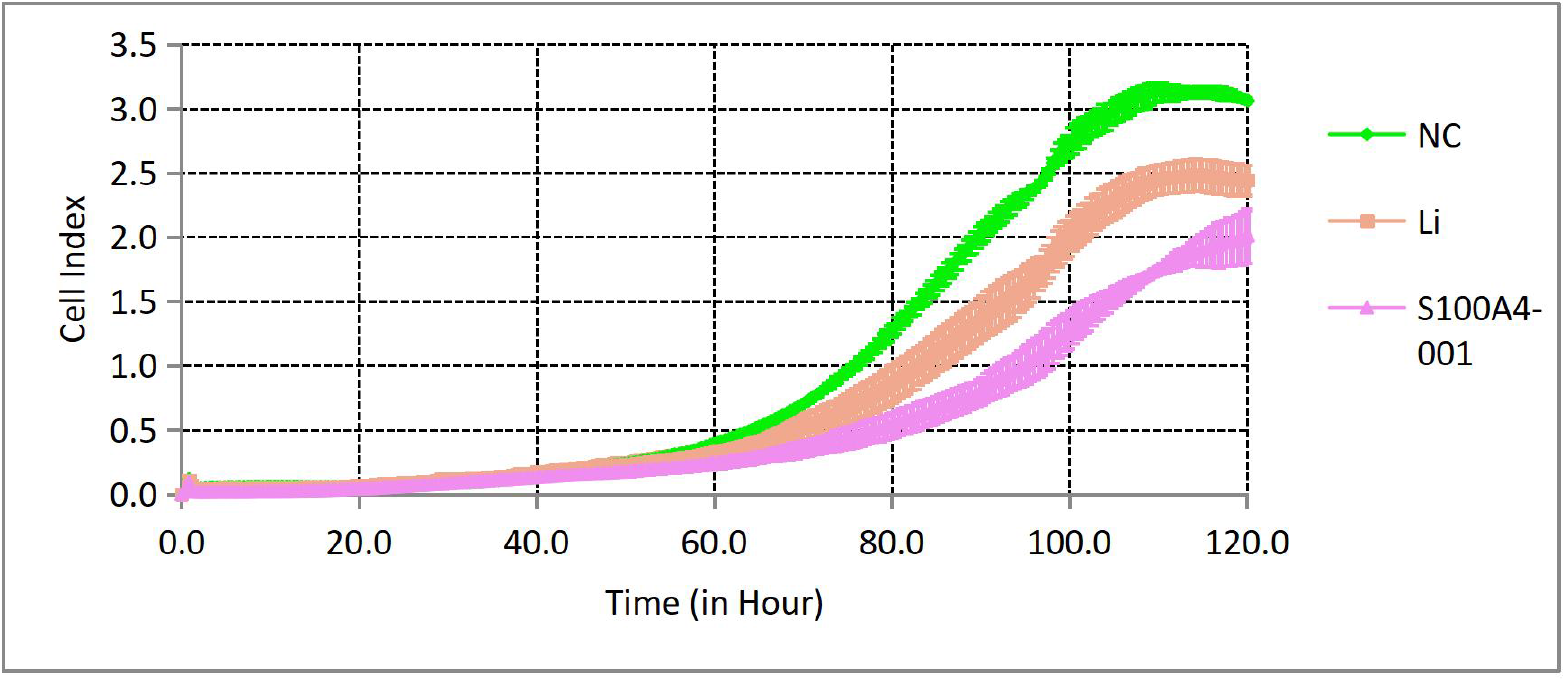
Real time cell analysis showing that knockdown of S100A4 inhibited cell migration of JAR cells

**Fig. 4.**
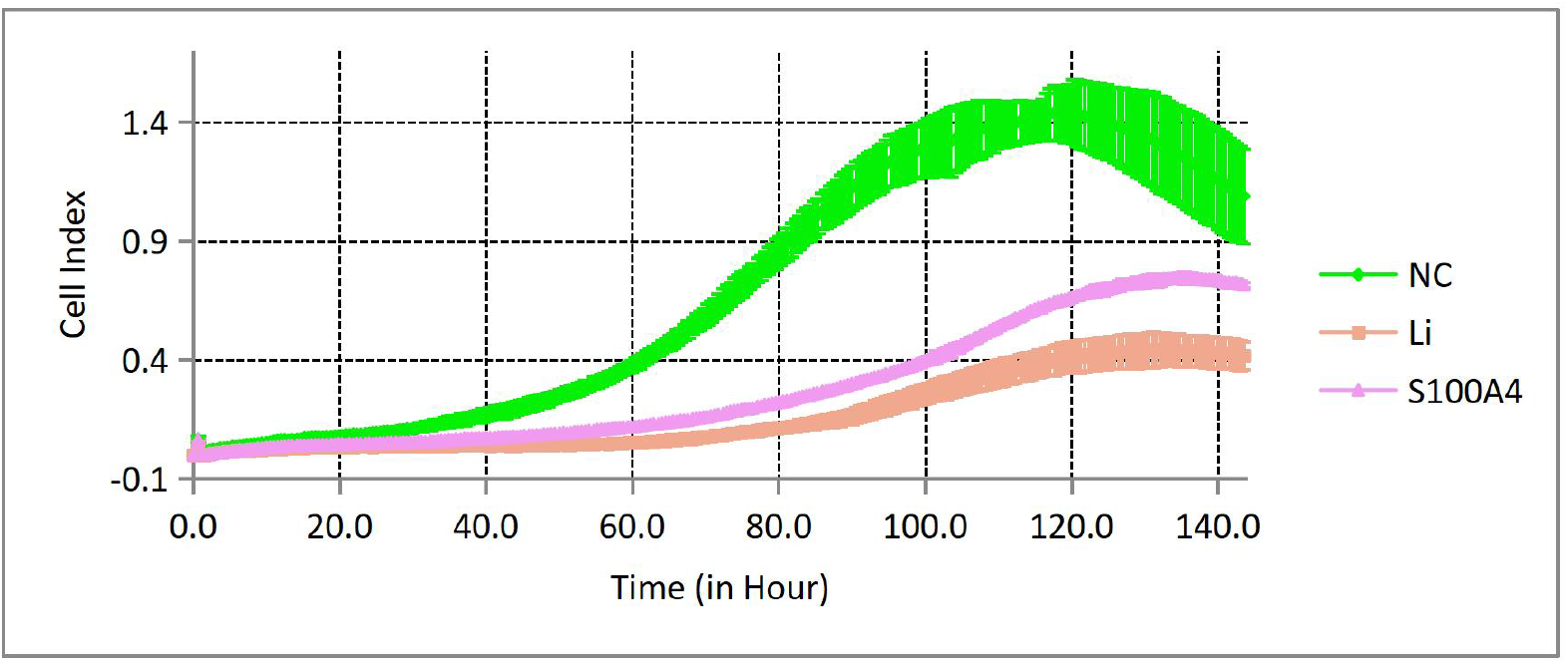
Real time cell analysis showing that knockdown of S100A4 did not reduce their invasion ability in JAR cells.

And then , the effect of S100A4 knockdown on apoptosis in JAR cells was investigated by flow cytometry and western blot, as shown in Figure 5. Both western blot (Fig. 5a) and flow cytometry(Fig. 5b) results demonstrated that S100A4 knockdown had no significant effect on apoptosis in JAR cells.

**Fig. 5.**
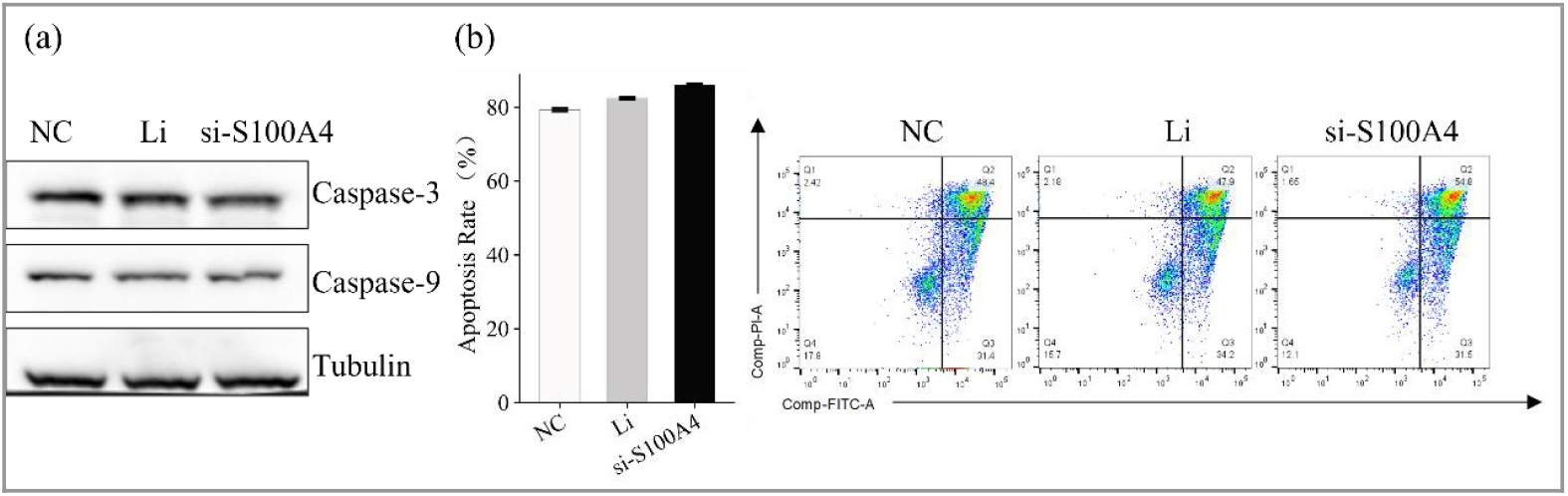
The apoptosis status of JAR cells following S100A4 knockdown. (a) Western blot analysis of apoptosis markers (caspase-3, caspase-9); (b) Flow cytometry analysis of Annexin V/PI staining. Both assays showing no significant effect of S100A4 knockdown on apoptosis.

Studies have demonstrated that S100A4 silencing promotes corneal burn wound healing through inhibition of the PI3K/Akt/mTOR pathway^15^. Additionally, S100A4 has been shown to be regulated by the PI3K/Akt signaling pathway in cancer^16^. The potential involvement of S100A4 in the altered biological behaviors of JAR cells might be associated with the PI3K/AKT signaling pathway. Therefore, we performed Western blot analysis to assess the expression status of key proteins in this pathway. Results showed that after knocking down the expression of S100A4 in JAR cells, IRS1 and PI3K protein expression was upregulated, whereas Akt1 expression was downregulated (Fig 6).

**Fig. 6.**
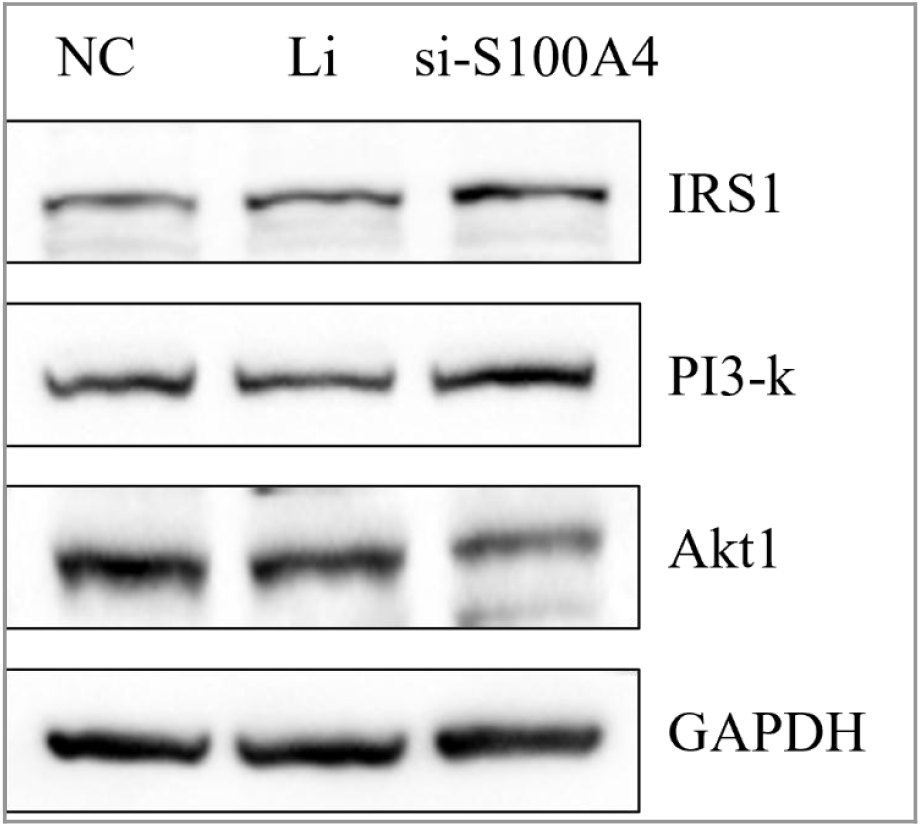
The expression of key proteins in the PI3K/AKT signaling pathway following S100A4 knockdown in JAR cells.

## 4. Discussion

This study suggests that S100A4 may exert phenotype-selective regulatory effects in choriocarcinoma. siRNA-mediated S100A4 knockdown significantly suppressed cellular proliferation and migration, but exhibited minimal impact on Matrigel invasion. Notably, both the lipofectamine control group and S100A4-knockdown group showed comparable invasion capacity, yet both were lower than the untreated cell control—indicating potential non-specific suppression from transfection reagents. This relative stability of the invasive phenotype contrasts with the canonical paradigm of S100A4 as a universal metastasis coordinator in epithelial cancers^17^. Notably, siRNA-mediated S100A4 knockdown did not induce significant apoptosis in JAR cells, as evidenced by concordant flow cytometry (Annexin V/PI staining) and Western blot data (unchanged cleaved caspase-3/9 levels).

The invasion resilience may relate to trophoblast-specific developmental programs. Physiological extravillous trophoblast invasion relies on mechanical deformation through vascular endothelia rather than protease-dependent matrix degradation^18^. The preserved transmigration capacity of knockdown cells through 8-μ m pores (simulating endothelial fenestrations) supports the existence of S100A4-bypass mechanisms in choriocarcinoma.Notably, S100A4, as a calcium-binding protein, directly interacts with non-myosin IIA (NMIIA) to regulate cellular motility and invasiveness^19^. However, the loss of S100A4 may be compensated through other mechanisms, such as maintaining cytoskeletal remodeling and cellular migratory capacity via the high expression of MYH9 and ITGB1. This compensatory mechanism may involve cytoskeletal reorganization and activation of signaling pathways, thereby sustaining normal cellular functions and adaptive responses.

Therapeutically, while targeting S100A4 effectively controls proliferation and migration phenotypes, its limited efficacy against invasion necessitates combinatorial approaches. In the context of choriocarcinoma, the inhibition of mechanotransduction pathways has shown promise in overcoming drug resistance and enhancing therapeutic efficacy. For example, the inhibition of brain-derived neurotrophic factor/tyrosine kinase B signaling suppresses choriocarcinoma cell growth, indicating the potential of targeting signaling pathways in this cancer type^20^. Additionally, metformin has been shown to regulate autophagy via LGMN to inhibit choriocarcinoma, further supporting the role of metabolic and mechanotransduction pathways in cancer therapy^21^. Mechanistically, S100A4 silencing triggered paradoxical signaling adaptations: IRS1 and PI3K expression increased, while Akt1 decreased and MEK1/2 remained unchanged (Fig. 6).

Limitations include: (1) Molecular evidence for compensatory pathways needs in vivo validation; (2) Clinical correlation between S100A4 expression and invasive phenotypes remains unestablished. Collectively, our data reveal a phenotype-signaling hierarchy: S100A4 silencing primarily disrupts proliferative/migratory programs via Akt1 suppression, while compensatory IRS1/PI3K elevation and MEK pathway stability preserve invasion. This functional decoupling underscores the need for combinatorial targeting (e.g., S100A4 inhibition + PI3K/mTOR blockade) to fully suppress choriocarcinoma dissemination.

